# Ligand-directed covalent labelling of a GPCR with a fluorescent tag

**DOI:** 10.1101/2020.04.21.053405

**Authors:** Leigh A Stoddart, Nicholas D Kindon, Omolade Otun, Clare R. Harwood, Foteini Patera, Dmitry B Veprintsev, Jeanette Woolard, Stephen J Briddon, Hester A Franks, Stephen J Hill, Barrie Kellam

## Abstract

Here, we describe rational design of a compound that covalently and selectively labels a G protein-coupled receptor (GPCR) in living cells with a fluorescent moiety. Using wild-type adenosine A_2A_ receptor as a model system, we show fluorescent labelling without impeding access to the orthosteric binding site and demonstrate its use in endogenously expressing systems. This offers a non-invasive and selective approach to study function and localisation of native GPCRs.

---

Fluorescent labelling a protein of interest can be achieved in many different ways and is an absolute prerequisite for many techniques used to probe both their function and localisation. For GPCRs, this is often achieved through use of protein tags (e.g. GFP, SNAP-tag), fluorescently labelled antibodies or fluorescent ligands^1^. Each of these technologies has their limitations and are not readily transferrable to endogenously expressing systems where GPCRs are expressed at low levels. Gene editing techniques, such as CRISPR/Cas9^2^, have allowed labelling of proteins^3^, including GPCRs^4,5^, at endogenous levels but the fluorophore used can dramatically alter expression levels^6^. An orthogonal approach to protein labelling uses ligand-directed chemistry whereby connecting a fluorophore via a highly reactive, electrophilic linker to a ligand that binds to the protein of interest. Upon binding of this conjugate, the linker can undergo a substitution reaction with a nucleophilic amino acid side chain (Lys, Ser, Tyr) close to the binding site, forming a new covalent bond between the fluorophore and protein (Figure 1a)^7^. The reactive groups used thus far for labelling proteins are bulky and not readily accommodated within the binding sites of a GPCR or suffer from slow reactivity^8,9^.

**Figure 1.**
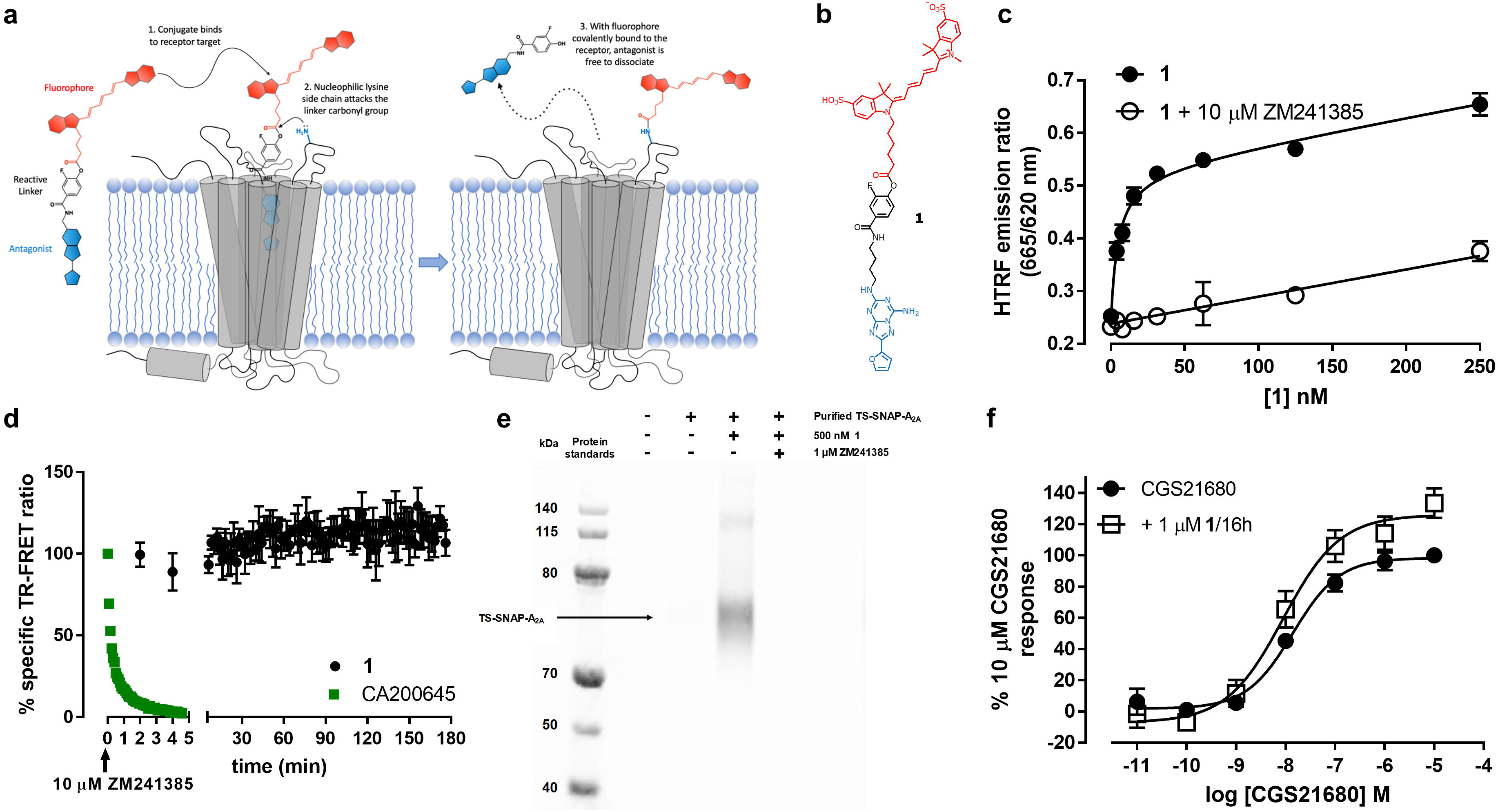
Design, pharmacological and biochemical characterisation of a 1. **(a)** Schematic illustration of ligand-directed labelling of A_2A_R by **1 (b)** Chemical structure of **1 (c)** TR-FRET saturation binding curves obtained by treating membranes containing Lumi4-Tb labelled SNAP-A_2A_R with increasing concentrations of **1** in the absence (closed circles) or presence (open circles) of 10μM ZM241385 for 1h at 37oC prior to determination of TR-FRET ratio. Data shown are representative of four experiments and each data point represents mean ± s.e.m of triplicate determinations. **(d)** Membranes containing Lumi4-Tb labelled SNAP-A_2A_R were pre-treated with 250nM **1**(5h, black circles) or CA200645 (2h, green squares) at 37°C prior to measurement of TR-FRET ratio. After basal reads, 10 μM ZM241385 was added and measurements taken every 2 min (**1**) or 5 sec (CA200645) for 5 min (CA200645) or 180 min (**1**). Non-specific binding was determined in the presence of 10μM ZM241385 and data normalized to total and non-specific binding at the zero time point. Each data point represents mean ± s.e.m. of four experiments each performed in triplicate. **(e)** T-Rex™-293 cells induced to express TS-SNAP-A_2A_R were treated with 500nM **1** in the presence or absence of 1μM ZM241385. Untreated cells were used as a control. TS-SNAP-A_2A_R was purified and separated on an SDS-PAGE gel. Direct Cy5 fluorescence was visualised using in-gel fluorescence. Gel shown is representative of three independent experiments. **(f)** CHO CRE-SPAP cells were treated with (open squares) or without (closed circles) 1μM **1** for 16h. Cells were washed for 30 min prior to the addition of increasing concentrations of CGS21680 and levels of CRE-mediated SPAP production measured after 5h. Data are normalized to basal (in the absence of agonist and **1**) and maximal CGS21680 response in the absence of **1** and each point represents the mean ± s.e.m. of five experiments performed in triplicate.

To overcome these difficulties, we incorporated a smaller phenyl ester group as the reactive moiety between the orthosteric head-group and fluorophore of a fluorescent conjugate for the adenosine A_2A_ receptor (A_2A_R), with the phenyl ring forming part of the binding pharmacophore. To enhance the chemical reactivity of the ester, molecular modelling suggested that the relatively small fluorine atom, the most electronegative element, could be introduced onto this phenyl ring and was compatible with binding of the antagonist into the binding site. If required, a further increase in chemical reactivity was envisaged by incorporating more fluorine atoms onto the phenyl ring. Based on recent fluorescent antagonists^10^ and the selective A__2A__R antagonist ZM241385^11^ we designed such a ligand, **1** (Figure 1b, Supplementary Figure 1), and used molecular modelling to guide close proximity of the phenyl ester with a nucleophilic amino acid side chain to allow transfer of a Cy5 fluorophore to the receptor (Figure 1a,b Supplementary Figure 2). To test its labelling efficacy in an over-expression system, we added increasing concentrations of **1** to membranes prepared from HEK293 cells expressing SNAP-A_2A_R labelled with Lumi4-Tb and measured time-resolved fluorescence resonance energy transfer (TR-FRET) between Lumi4-Tb and the Cy5 subsequently attached to the receptor by **1** via ligand-directed covalent labelling. We detected a concentration-dependent increase in TR-FRET ratio which could be inhibited by co-incubation with the unlabelled competitive antagonist ZM241385 (10μM; Figure 1c). This was consistent with specific binding to the A_2A_R and resulted in an estimated equilibrium dissociation constant (K_D_) of 5.5±2.0nM for **1** (n = 4).

To test the irreversible nature of this binding interaction, we added an excess of ZM241385 (10μM) to membranes previously labelled with 250nM **1** and observed very little change in the TR-FRET ratio over the subsequent 3 h (103.2±6.8% vs 0 time point, n=4). This suggested that the fluorophore remained in close proximity to the Lumi4-Tb labelled receptor and could not be displaced by ZM241385, indicating either permanent transfer of the flurophore or very slow dissociation of the ligand from the receptor (Figure 1d). In marked contrast, when we undertook a similar experimental protocol with the competitive orthosteric fluorescent antagonist CA200645, the TR-FRET ratio returned to baseline within five minutes of addition of ZM241385 (Figure 1d, Supplementary Figure 3) indicating full dissociation from the receptor. To confirm covalent labelling of A_2A_R by **1**, T-Rex™-293 cells expressing a Twin-Strep-SNAP-A_2A_R (TS-SNAP-A_2A_R) construct were treated with **1** prior to purification, separation by SDS-PAGE and visualisation of labelled samples by in-gel fluorescence. A strong band corresponding to Cy5 labelled TS-SNAP-A_2A_R was observed at 73 kDa, the expected molecular weight of a monomer, and a concomitant weaker band at 130 kDa (Figure 1e, Supplementary Figure 4). Labelling of A_2A_R with Cy5 was prevented when the cells were co-incubated with **1** and ZM241385 (1μM). The presence of Cy5 fluorescence after purification and denaturation of A_2A_R confirms the hypothesis that a covalent bond is present between receptor and fluorophore.

**1** was designed to label the A_2A_R with Cy5 and then allow dissociation of the pharmacophore portion of the ligand from the orthosteric binding site, enabling additional ligands to subsequently bind to the receptor. However, it is conceivable that labelling the receptor with Cy5 close to the entrance of the binding pocket may act as a structural barrier and prevent access of additional ligands^12^. The A_2A_R is a G_s_ coupled receptor and its activation leads to an increase in cAMP and subsequent transcription of genes under the control of cAMP response element (CRE). Therefore, to determine whether the orthosteric binding site was accessible after traceless Cy5 labelling of the receptor, we evaluated the ability of the A_2A_-selective agonist CGS21680 to stimulate CRE-mediated gene expression in CHO CRE-SPAP (secreted placental alkaline phosphatase) cells expressing the A_2A_R. In naïve cells, increasing concentrations of CGS21680 produced a concentration-dependent increase in SPAP production (pEC_50_ = 7.86±0.09, *n* = 5). When cells were treated overnight (16h) with 1μM 1, conditions which lead to a significant fluorescent labelling of the A_2A_R, and washed prior to addition of the agonist, CGS21680 remained equally potent compared to untreated cells (pEC_50_ = 8.05±0.16, *n* = 5; *p* = 0.32, unpaired t-test). A small increase in the maximum response (13.3 ± 5.2% increase in maximal SPAP production vs control, Figure 1f) is unlikely to be biologically significant. These data demonstrate that following covalent labelling of the receptor by **1** the antagonist pharmacophore readily dissociates from the orthosteric receptor binding site. Furthermore, agonist access to the orthosteric binding site was not impaired by the covalent attachment of Cy5 to the receptor.

We visualised the Cy5 labelling of SNAP-A_2A_R by **1** in live HEK293 cells using confocal microscopy. We observed clear membrane localisation of the Cy5 fluorescence in the continued presence of **1** and a high degree of co-localisation with the AF488 labelled SNAP-A_2A_R (Figure 2a). This fluorescence labelling was prevented in the presence of ZM241385 (10μM), demonstrating again that the ligand needs to bind to the orthosteric binding site of the A_2A_R in order to transfer the fluorescent label. After labelling with 250nM **1** for 2h, we observed no change in fluorescence intensity at the cell surface despite washing cells four times over 1h (Figure 2a-c). The addition of a high concentration of ZM241385 (10μM) after pre-labelling with **1** also had no influence on the level of Cy5 fluorescence (Figure 2b, Supplementary Figure 5). These data demonstrate that **1** can label A_2A_R irreversibly with Cy5 in a live-cell system.

**Figure 2.**
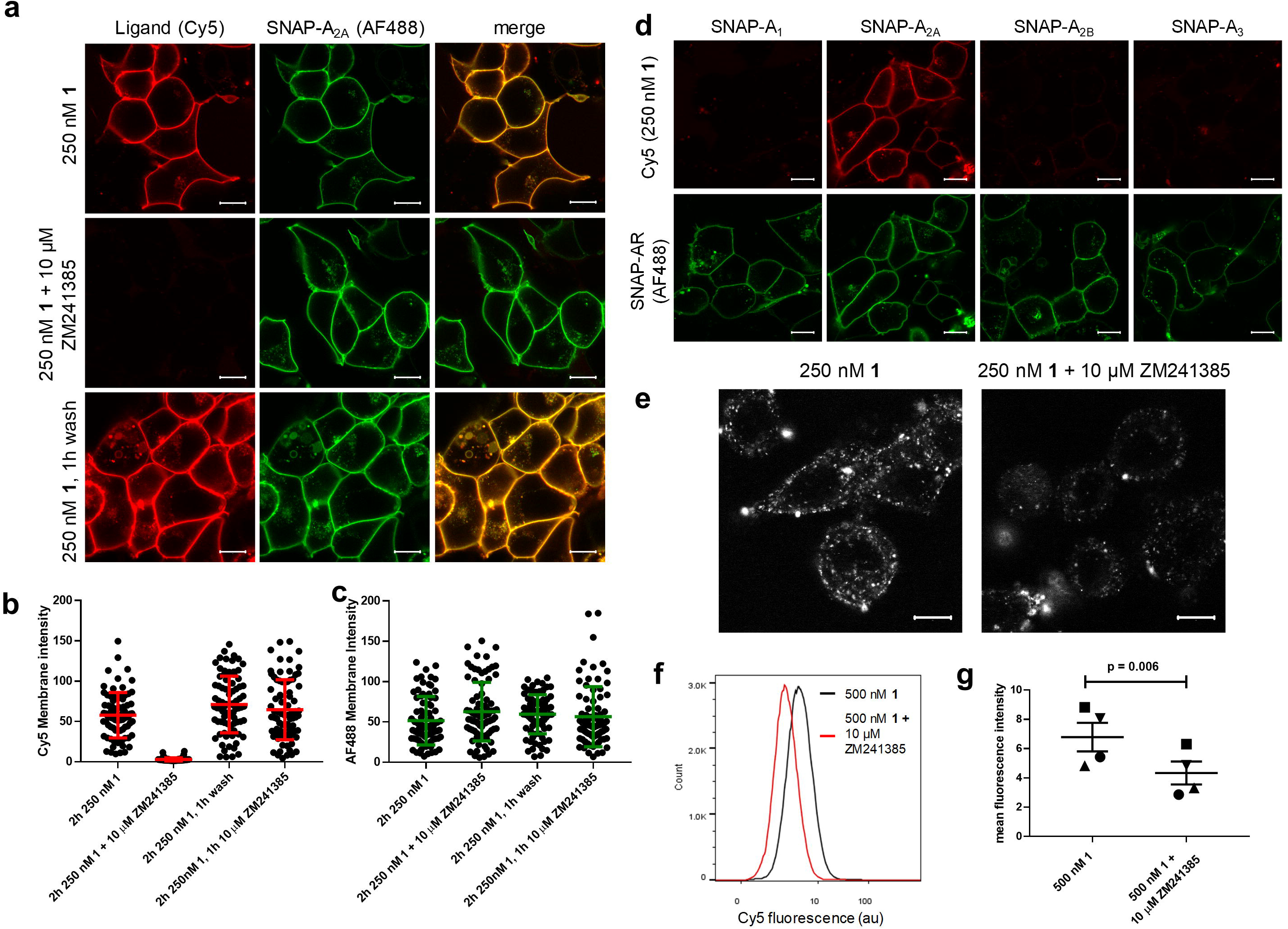
1 is selective and can be used to visualise A_2A_R in live cells and endogenously expressing systems. **(a)** Live HEK293 cells expressing SNAP-A_2A_R were labelled with SNAP-surface-AF488 and then treated with 250nM **1** in the absence (top row) or presence (middle row) of ZM241385 for 2h prior to the capture of single equatorial confocal images. Cells treated with 250nM **1** were then washed multiple times over 1h and then imaged (bottom row). Left hand column represents Cy5 fluorescence, middle column AF488 fluorescence and right hand column the merged image. Fluorescent intensity values were determined for Cy5 **(b)** and AF488 **(c)** in membrane regions of interest from the images obtained as described in **a** and also in cells treated with 10μM ZM241385 for 1h after 2h incubation with 250 nM **1**(images shown in supplementary figure 4). Each point represents the values obtained from one cell. Images were obtained in four independent experiments, error bars represent mean ± SD. **(d)** HEK293 cells expressing SNAP tagged versions of one of the four adenosine receptor subtypes (A_1_, A_2A_, A_2B_ or A_3_) were labelled with SNAP-surface-AF488 prior to the addition of 250nM **1** for 2h prior to the capture of single equatorial images. **(e)** SK-BR-3 cells were labelled with 250nM **1** in the presence (right hand image) or absence (left hand image) of 10μM ZM241385 for 2h prior to the capture of single equatorial confocal images. Images shown in **a, d** and **e** are representative of images taken in four (**a**) or three (**d** and **e**) independent experiments, with all image sets taken using identical settings for laser power, gain, and offset in both channels. Scale bar shown represents 10 μm. **(f)** Representative flow cytometry histograph of human macrophages treated with 500nM **1** (black line) or 500nM **1** plus 10μM ZM241385 (red line) for 2h at 37°C. **(g)** Mean fluorescence intensity of macrophages derived from four healthy donors treated as in **(f)** (p = 0.006, paired t-test). Each symbol represents one donor.

One of the advantages of ligand-directed labelling of proteins over other labelling methodologies is that selectivity between closely related proteins can be achieved through the use of a ligand that binds selectively to the protein of interest. To check the selectivity of **1** itself, we treated HEK293 cells expressing the three other adenosine receptor subtypes (A_1_R, A_2B_R, A_3_R) with **1** and determined the levels of labelling with Cy5 using confocal imaging (Figure 2d). We observed no specific Cy5 fluorescence in cells expressing A_1_R, A_2B_R or A_3_R. This high degree of selectivity of **1** for A_2A_R over the other three adenosine receptor subtypes cannot be attributed solely to the selectivity of ZM241385 as it is reported to be only 60 fold selective for A_2A_R over A_2B_R^11^. It is therefore likely that this selectivity also results from close proximity of nucleophilic residues only within A_2A_R to the reactive core of **1** when it is resident within the orthosteric binding site of the receptor (Supplementary Figure 6).

As **1** showed selectivity for A_2A_R over other adenosine receptor subtypes, we then investigated if it could be used to label endogenously expressed A_2A_Rs. For this we selected the human breast cancer cell line SK-BR-3 and human monocyte-derived macrophages which are both known to express the A_2A_R^13–15^. Live-cell confocal imaging of SK-BR-3 cells treated with **1** showed clear cell surface Cy5 labelling, which was prevented by the presence of ZM241385 (Figure 2e) and ZM241385-sentitive labelling of macrophages with Cy5 was observed via flow cytometry (Figure 2f, p<0.0001, two-sided unpaired T-test with Welch’s correction; Figure 2g, p=0.006, two-sided paired T-test; Supplementary Figure 7) demonstrating labelling of endogenous A_2A_R with **1**.

In summary, we have described rational design of a compound that can covalently and selectively label a GPCR with a florescent molecule without affecting the binding site of the receptor. Ligand directed labelling of GPCRs is a novel, non-invasive approach to visualise GPCRs and opens up the possibilities to study ligand binding, receptor trafficking and signalling in endogenously and clinically relevant systems.

## Acknowledgements

The authors thank the School of Life Sciences imaging facility (SLIM), University of Nottingham for their contribution to the imaging in this publication and Dr Mark Soave and Dr Bradley Hoare (University of Nottingham) for help in generating the constructs used in this study. This project was funded the Medical Research Council [grant number MR/N020081/1] awarded to J.W., S.J.B, S.J.H and B.K. O.O was supported by a COMPARE Vacation Studentship.

## Author Contributions

L.A.S., N.D.K., O.O., C.H., F.P. and H.A.F. performed the research. L.A.S., N.D.K., S.J.B., H.A.F., S.J.H. and B.K. designed the research study. L.A.S., N.D.K., O.O., C.H., F.P. and H.A.F analysed the data. L.A.S., N.D.K., J.W., D.B.V., S.J.B, S.J.H. and B.K. wrote the paper.

