## Supplementary Information and Figures for "Ligand-directed covalent labelling of a GPCR with a fluorescent tag"

### Methods

#### Chemistry

Chemicals and solvents of analytical and HPLC grade were purchased from commercial suppliers and used without further purification. Sulfo-Cyanine5 carboxylic acid was purchased from Lumiprobe. Reactions were monitored by thin-layer chromatography on commercially available silica pre-coated aluminium-backed plates (Merck Kieselgel 60 F254). Visualisation was under UV light (254 nm and 366 nm). Flash column chromatography was performed using silica gel 60, 230-400 mesh particle size (Sigma Aldrich). NMR spectra were recorded on a Bruker-AV 400. $^{1}$H NMR spectra were recorded at 400.13 MHz and $^{13}$C NMR spectra at 126.00 MHz. Chemical shifts (δ) are recorded in parts per million (ppm) and coupling constants are recorded in Hz. The following abbreviations are used to described signal shapes and multiplicities; singlet (s), doublet (d), triplet (t), quadruplet (q), broad (br), dd (doublet of doublets), ddd (double doublet of doublets), dtd (double triplet of doublets) and multiplet (m). Processing of the NMR data was carried out using the NMR software Topspin. LC-MS spectra were recorded on a Shimadzu UFLCXR system coupled to an Applied Biosystems API2000 and visualised at 254 nm (channel 1) and 220 nm (channel 2). LC-MS was carried out using a Phenomenex Gemini-NX C18 110A, column (50 mm × 2 mm x 3 μm) at a flow rate 0.5 mL/min over a 5 min period. All high resolution mass spectra (HRMS) were recorded on a Bruker microTOF mass spectrometer using MS electrospray ionization operating in positive or negative ion mode. RP-HPLC was performed on a Waters 515 LC system and monitored using a Waters 996 photodiode array detector at wavelengths between 190 and 800 nm. Spectra were analysed using Millenium 32 software. Semi-preparative HPLC was performed using YMC-Pack C8 column (150 mm × 10 mm × 5 μm) at a flow rate of 5.0 mL/min using a gradient method of 20-70% B over 15 minutes (Solvent A = 0.1%

formic acid in $H_2O$, solvent B = 0.1% formic acid in $CH_3CN$). Analytical RP-HPLC was performed using a YMC-Pack C8 column (150 mm × 4.6 mm × 5 μm) at a flow rate of 1.0 mL/min. Final products were one single peak and >98% pure. The retention time of the final product is reported using a gradient method of 10-95% solvent B in solvent A over 26 min. (Solvent A = 0.1% formic acid in $H_2O$, solvent B = 0.1% formic acid in $CH_3CN$).

**2-(Furan-2-carbonyl)hydrazine-1-carboximidamide (3)[1]**: 2-Furoic hydrazide (**2**) (6.0 g, 47.5 mmol) and 2-methyl-2-thiopseudourea hemisulfate salt (33.1 g, 238 mmol) were added to a stirred solution of sodium hydroxide (3.04 g, 76 mmol) in water (150 ml) at room temperature. After 24h, the precipitate was collected by filtration and washed with water and diethyl ether to give 2-(furan-2-carbonyl)hydrazine-1-carboximidamide. Yield 4.00 g (23.8 mmol, 50%) and used directly in the next step without further purification.

**3-(Furan-2-yl)-1*H*-1,2,4-triazol-5-amine (4)[1]**: 2-(Furan-2-carbonyl)hydrazine-1-carboximidamide (**3**) (4.0 g, 23.8 mmol) was suspended in water (70 ml) and the mixture heated under reflux for 24 h. The reddish solution was evaporated to dryness and the resulting solid slurried in water, collected by filtration and dried to give 3-(furan-2-yl)-1*H*-1,2,4-triazol-5-amine. Yield 2.8 g (18.6 mmol, 78%). $^1$H NMR (400 MHz, DMSO) δ 12.07 (s, 1H), 7.68 (s, 1H), 6.68 (d, $J$ = 3.3 Hz, 1H), 6.54 (s, 1H), 6.07 (s, 2H).

**2-(Furan-2-yl)-5-(methylthio)-[1,2,4]triazolo[1,5-*a*][1,3,5]triazin-7-amine (5)[1]**: Dimethyl *N*-cyanodithioiminocarbonate (2.2 g, 15 mmol) and 3-(furan-2-yl)-1*H*-1,2,4-triazol-5-amine (2.2 g, 14.6 mmol) were mixed and heated at 170$^o$C under nitrogen for 1 h. After cooling the crude reaction mixture was dissolved in methanol and dichloromethane and absorbed on to silica. Purification was by silica gel chromatography eluting with 5-10 % ethyl acetate in dichloromethane to give 2-(furan-2-yl)-5-(methylthio)-[1,2,4]triazolo[1,5-*a*][1,3,5]triazin-7-amine. Yield 1.4 g (5.6 mmol, 38%). LC/MS (m/z) 248.9 (M+1), r.t. 2.47 min, $^1$H NMR (400 MHz, DMSO) δ 9.11 – 8.54 (br s,

2H), 7.93 (dd, *J* = 1.8, 0.8 Hz, 1H), 7.17 (dd, *J* = 3.4, 0.8 Hz, 1H), 6.72 (dd, *J* = 3.4, 1.8 Hz, 1H), 2.52 (s, 3H).

**2-(Furan-2-yl)-5-(methylsulfonyl)-[1,2,4]triazolo[1,5-*a*][1,3,5]triazin-7-amine (6)[1]**: 3-chloroperbenzoic acid (3.8 g, 22 mmol) was added portionwise over 3 minutes to a stirred suspension of 2-(furan-2-yl)-5-(methylthio)-[1,2,4]triazolo[1,5-*a*][1,3,5]triazin-7-amine (1 g, 4.0 mmol) in dichloromethane (100 ml). Initially a clear solution formed and then a precipitate. After 16 h, the mixture was concentrated to 10 ml and ethanol (10 ml) added. The mixture was concentrated to 10 ml and the product allowed to crystallise out over 3 h which was collected by filtration to give 2-(furan-2-yl)-5-(methylsulfonyl)-[1,2,4]triazolo[1,5-*a*][1,3,5]triazin-7-amine. Yield 0.9 g (3.2 mmol, 80%). LC/MS (m/z) 281.2 (M+1), r.t. 2.12 min, $^{1}$H NMR (400 MHz, DMSO) δ 9.81 (s, 1H), 9.48 (s, 1H), 7.99 (dd, *J* = 1.8, 0.8 Hz, 1H), 7.27 (dd, *J* = 3.4, 0.8 Hz, 1H), 6.76 (dd, *J* = 3.5, 1.8 Hz, 1H), 3.37 (s, 3H).

**2-(Furan-2-yl)-[1,2,4]triazolo[1,5-*a*][1,3,5]triazin-5-yl)amino)butyl)carbamate (7)**: *tert*-Butyl *N*-(3-aminobutyl)carbamate (0.3 g, 1.6 mmol) in acetonitrile (1.5 ml) was added to a stirred solution of 2-(furan-2-yl)-5-(methylsulfonyl)-[1,2,4]triazolo[1,5-*a*][1,3,5]triazin-7-amine(0.3 g, 1.07 mmol) in acetonitrile (3 ml). After 2h, the reaction mixture was absorbed onto isolute and applied to a silica gel column eluting with 5-10% methanol in dichlormethane to give tert-butyl (4-((7-amino-2-(furan-2-yl)-[1,2,4]triazolo[1,5-*a*][1,3,5]triazin-5-yl)amino)butyl)carbamate. Yield 0.3 g (0.78 mmol, 73%). $^{1}$H NMR (400 MHz, DMSO) δ 8.25 – 7.93 (m, 2H), 7.87 (d, *J* = 1.8 Hz, 1H), 7.54 – 7.35 (m, 1H), 7.04 (d, *J* = 3.6 Hz, 1H), 6.82 – 6.77 (m, 1H), 6.68 (dd, *J* = 3.5, 1.9 Hz, 1H), 3.30 – 3.19 (m, 2H), 2.93 (q, *J* = 6.5 Hz, 2H), 1.53 – 1.46 (m, 2H), 1.45 – 1.39 (m, 2H), 1.37 (s, 9H)

***N*-(4-((7-amino-2-(furan-2-yl)-[1,2,4]triazolo[1,5-*a*][1,3,5]triazin-5-yl)amino)butyl)-3-fluoro-4-hydroxybenzamide (8)**: *tert*-Butyl (4-((7-amino-2-

(furan-2-yl)-[1,2,4]triazolo[1,5-*a*][1,3,5]triazin-5-yl)amino)butyl)carbamate (0.035 g, 0.09 mmol) in dichloromethane (5 ml) was treated with trifluoroacetic acid (2 ml). After 35 minutes the solution was evaporated to dryness. Toluene (10 ml) was added and the mixture evaporated to dryness. This process was repeated 2 more times to remove residual trifluoracetic acid to give $N^5$-(4-aminobutyl)-2-(furan-2-yl)-[1,2,4]triazolo[1,5-*a*][1,3,5]triazine-5,7-diamine trifluoroacetate salt which was used without further purification.

1-[Bis(dimethylamino)methylene]-1*H*-1,2,3-triazolo[4,5-*b*]pyridinium 3-oxide hexafluorophosphate (HATU) (0.041 g, 0.1 mmol) was added to a stirred solution of 3-fluoro-4-hydroxybenzoic acid (0.014 g, 0.09 mmol) and diisopropylethylamine (0.15 ml) in DMF (1 ml). After 0.5h, the slurry was added to the $N^5$-(4-aminobutyl)-2-(furan-2-yl)-[1,2,4]triazolo[1,5-*a*][1,3,5]triazine-5,7-diamine trifluoroacetate salt and diisopropylethylamine (0.25 ml) in DMF (1 ml). The mixture was heated at 90 $^oC$ for 2h. After cooling, the solvent was removed under high vacuum. Purification was by silica gel chromatography eluting with 5%-10% methanol in dichloromethane to give *N*-(4-((7-amino-2-(furan-2-yl)-[1,2,4]triazolo[1,5-*a*][1,3,5]triazin-5-yl)amino)butyl)-3-fluoro-4-hydroxybenzamide. Yield 0.03 g (0.07 mmol, 70%) as a white solid. LC/MS (m/z) 427.6 (M+1), r.t. 2.31 min, $^1$H NMR (400 MHz, DMSO) δ 10.42 (s, 1H), 8.30 (t, *J* = 5.6 Hz, 1H), 8.20 – 8.00 (br s, 2H), 7.87 (d, *J* = 1.8 Hz, 1H), 7.63 (dd, *J* = 12.3, 2.1 Hz, 1H), 7.55 (dd, *J* = 8.4, 2.1 Hz, 1H), 7.55 – 7.40 (m, 1H), 7.04 (d, *J* = 3.5 Hz, 1H), 6.98 (t, *J* = 8.6 Hz, 1H), 6.68 (dd, *J* = 3.5, 1.8 Hz, 1H), 3.32 – 3.20 (m, 4H), 1.59 – 1.53 (m, 4H). $^{13}$C NMR (126 MHz, DMSO) δ 165.19, 161.65, 159.68, 156.27, 150.81 (d, J 240Hz) 150.44, 148.09 (d, J 11.4Hz), 146.70, 145.08, 126.39 (d, J 5.4Hz), 124.60 (d, J 2.52Hz), 117.55 (d, 3.78Hz), 115.57 (d, J 18.9Hz), 112.37, 112.05, 27.20, 26.88. (2 x $CH_2$ under DMSO peak)

**2-((1*E*,3*E*)-5-((*E*)-1-(6-(4-((4-((7-Amino-2-(furan-2-yl)-[1,2,4]triazolo[1,5-*a*][1,3,5]triazin-5-yl)amino)butyl)carbamoyl)-2-fluorophenoxy)-6-oxohexyl)-**

**3,3-dimethyl-5-sulfoindolin-2-ylidene)penta-1,3-dien-1-yl)-1,3,3-trimethyl-3*H*-indol-1-ium-5-sulfonate (1)**: 2-Bromo-1-ethyl-pyridinium tetrafluoroborate (BEP reagent)(0.32 mg, 1.17x10-6 mol) and diisopropylethylamine (10mg, 7.75x10-5 mol) in dry *N*,*N*-dimethylformamide (0.6 ml) was added to sulfo-cyanine-C5 carboxylic acid (1 mg, 1.47x10$^{-6}$ mol). After 15 minutes in the dark, this solution was added to *N*-(4-((7-amino-2-(furan-2-yl)-[1,2,4]triazolo[1,5-a][1,3,5]triazin-5-yl)amino)butyl)-3-fluoro-4-hydroxybenzamide (**8**) (0.7 mg, 1.6x10$^{-6}$ mol). After 16 h in the dark, the solution was evaporated to dryness under high vacuum. Purification was by semi-preparative RP-HPLC method B. The product containing fractions were combined and concentrated to remove most of the acetonitrile then lyophilised to give 2-((1*E*,3*E*)-5-((*E*)-1-(6-(4-((4-((7-amino-2-(furan-2-yl)-[1,2,4]triazolo[1,5-*a*][1,3,5]triazin-5-yl)amino)butyl)carbamoyl)-2-fluorophenoxy)-6-oxohexyl)-3,3-dimethyl-5-sulfoindolin-2-ylidene)penta-1,3-dien-1-yl)-1,3,3-trimethyl-3*H*-indol-1-ium-5-sulfonate as a blue solid. Yield 1.0 mg (9.5x10-7 mol, 65%). HMRS (m/z) M-H calculated for $C_{51}H_{55}FN_{10}O_{10}S_2$: 1049.3455, found M-H: 1049.3452. Analytical RP-HPLC, retention time 13.80 minutes.

**cDNA constructs** We generated SNAP-labelled adenosine receptor constructs by amplifying the full length sequence of SNAP-tag (New England Biolabs, Ipswich, MA) and fusing it in frame with the membrane signal sequence of the $5HT_{3A}$ receptor with pcDNA3.1 to yield sig.SNAP. We then fused the full-length human sequence of each of the four adenosine receptor subtypes (with the methionine start signal removed) to the 3′ end of the sig.SNAP in pcDNA3.1. This gave the constructs designated as SNAP-$A_1$R, SNAP-$A_{2A}$R, SNAP-$A_{2B}$R and SNAP-$A_3$R, all of which contain the signal sequence. We generated the pcDNA4TO-TS-SNAP-$A_{2A}$ construct by amplifying the $A_{2A}$ receptor (with the methionine start signal removed) and inserting into a pcDNA4TO vector (ThermFisher Scientific, Paisley, UK) containing a Twin-Step-tag® sequence and the SNAP sequence using Gibson assembly[2]. This gave the construct designated as TS-SNAP-$A_{2A}$R.

**Ligands.** CA200645 was from HelloBio (Bristol, UK). ZM 241385 and CGS21680 were from Tocris (Bristol, UK).

**Cell culture and transient transfection.** Human embryonic kidney 293 (HEK293) cells expressing the GloSensor cAMP biosensor (HEKG) were obtained from Promega (Southampton, UK) and were maintained in Dulbecco's modified Eagle's medium (DMEM) containing 10% fetal calf serum (FCS) and L-glutamine. T-REx$^{TM}$-293 cells (Invitrogen) stably expressing the pcDNA4TO TwinStrepSNAP-$A_{2A}$ (TS-SNAP-$A_{2A}$) construct were maintained in high glucose DMEM containing 10% FCS, 5 µg/µL blasticidin and 20 µg/µL zeomycin. SNAP-$A_{2A}$ HEKG cells were generated as previously described[3]. Chinese hamster ovary (CHO) cells stably expressing a cAMP response element-secreted placental alkaline phosphatase (CRE-SPAP) reporter gene expressing the human $A_{2A}$R were generated as described previously[4] and maintained in Dulbecco's Modified Eagle Medium:Nutrient Mixture F-12 (DMEM/F12) medium containing 10% FCS and 2 mM L-glutamine. SK-BR-3 cells obtained from ATCC (Manassas, VA) were maintained in McCoy's 5a Medium supplemented with 10% FCS. All cell lines were maintained at 37$^{o}$C in a humidified atmosphere of air/$CO_2$ (19:1). For transient transfections, HEK293G cells grown in 8-well chamber slides were transfected with the required SNAP-adenosine receptor construct using Fugene (Promega) according to the manufacturers' instructions 24h prior to experiment.

**Human macrophage generation.** Heparinised whole blood was obtained by venepuncture from the antecubital fossa of the arm of healthy volunteers after written informed consent (Ethics from University of Nottingham Ethics committee, ref 161-1711). Peripheral blood mononuclear cells (PBMC) were immediately separated by density centrifugation over Ficoll Histopaque 1077 (Sigma) followed by washes in endotoxin-free phosphate-buffered saline (PBS, Sigma). PBMC were washed in MACS buffer (PBS + 1% fetal calf serum (FCS, Sigma) + 2µM EDTA (Sigma)) then incubated with CD14 microbeads (Miltenyi Biotech) and monocytes isolated by magnetic separation (purity routinely >95%). Purified CD14+ monocytes were differentiated into macrophages at 37°C/5%$CO_2$ for 7 days at 1x$10^6$/well in low-attachment 24-well plates (Corning Costar) in macrophage medium (RPMI 1640 (Sigma) supplemented with 10% FCS and 1% sodium pyruvate (Sigma)) and 20U/ml recombinant human granulocyte-macrophage colony-stimulating factor (Peprotech) with medium plus additives supplemented at day 4.

**Macrophage 1 labelling and flow cytometry.** Medium was aspirated from day 7 macrophages and 500 nM **1** in the presence or absence of 10 μM ZM241385 added in fresh macrophage medium whilst cells remained in the differentiation plate. Cells were incubated for 2 hours at 37°C/5% $CO_2$ then medium aspirated, wells washed with cold PBS and macrophages harvested by incubating on ice in cold PBS for 25 minutes. Cy5 fluorescence was measured by flow cytometry (MacsQuant, Miltenyi Biotech) and analysed using FlowJo software. Gating strategy is demonstrated in Supplementary Figure 7.

**Lumi4-Tb labelling and membrane preparation.** SNAP-$A_{2A}$ HEKG cells were grown to confluence in poly-D-lysine coated T175 flasks. Cells were washed once in PBS, 100 nM SNAP-Lumi4-Tb in 1x Lab Med (CisBio, Bagnols-sur-Ce'ze, France) was added and incubated for 1h at 37°C. After 1h, SNAP-Lumi4-Tb solution was removed and replaced with ice-cold PBS, and the cells were removed from the flask using a cell scraper. The cells were then transferred to a 50 ml tube and centrifuged at 250x*g* for 10 min. The supernatant was discarded and the resulting pellets were stored at -80°C. For membrane preparation, cell pellets were thawed and resuspended in ice-cold PBS and homogenised using an IKA T10 Ultra-Turrax disperser in 10 x 5 s bursts at 15,000 rpm. After removal of unbroken cells and nuclei by centrifugation at 1200xg for 10 min, the resulting supernatant was centrifuged at 41,415x*g* for 30 min to obtain the membrane pellet. The pellet was then resuspended in ice-cold PBS and homogenised by 20 passes at 1000 rpm using a Kartell serrated pestle and a borsilicate glass homogeniser mortar fitted to an IKA RW16 overhead stirrer. Protein concentration of the resuspended membranes was determined using a bicinchoninic acid protein assay and SNAP-Lumi4-Tb membranes stored at -80°C until required.

**TR-FRET binding assay.** All TR-FRET assays were performed in opaque bottomed 96-well plates and read on a PHERAstar FS (BMG Labtech,Offenberg, Germany) with the terbium (donor) excited with 30 flashes of laser at 337 nm and emission collected at 620 nm (terbium) and 665 nm (Cy5/BY630) 400 ms after excitation. The TR-FRET ratio was calculated by dividing the Cy5/BY630 emission (665 nM) by the terbium emission (620 nm. For membrane saturation TR-FRET binding assays, 2.5 mg of Lumi4-Tb labelled SNAP-$A_{2A}$ membranes were incubated with the required compounds in HEPES buffered saline solution (HBSS: 145 mmol/L NaCl, 5 mmol/L KCl, 1.7 mmol/L $CaCl_2$, 1 mmol/L $MgSO_4$, 10 mmol/L HEPES, 2 mmol/L sodium pyruvate, 1.5 mmol/L $NaHCO_3$, 10 mmol/L D-glucose, pH 7.4) containing 1 mg/ml saponin for 1 h at 37°C before reading on the PHERAstar. For dissociation experiments, 2.5 µg of SNAP-$A_{2A}$ membranes were incubated with compounds in HBSS plus saponin for 5 h for **1** and 2 h for CA200645 at 37°C in 96-well plates. After required incubation time, basal TR-FRET readings were taken on the PHERAstar, for **1** treated membranes 10 µM ZM241385 was added to each well manually in a 1:1 dilution to ensure adequate mixing and TR-FRET readings were then taken every 5 min for 3h as detailed above. For CA200645, 10µM ZM241385 was added using the inbuilt PHERAstar injectors. Due to the rapid dissociation of CA200645, readings were taken every 5 sec for 5 min with 20 flashes per read.

**Labelling of cells with 1 for purification and in gel fluorescence** TS-SNAP-$A_{2A}$ TREx-293 cells were grown to 70% confluence prior to induction of TS-SNAP-$A_{2A}$ expression by the addition of 1µg/mL tetracyclin to normal growth medium. After 50 h induction, medium was replaced and to the required flasks 500 nM **1** or 500 nM **1** plus 1 µM ZM241385 added and cells incubated for a further 5h at 37°C/5% $CO_2$. After 5h, medium was removed and cells washed twice with phosphate buffered saline (PBS). Cells were then detached from flasks using cell dissociation solution non-enzymatic (Sigma), washed off with PBS and solutions removed from the flasks. Cell suspensions were spun at 362 *x g* for 10mins. Supernatant was aspirated and cell pellets frozen at -80°C until use.

**Solubilisation and purification of 1 labelled TS-SNAP-$A_{2A}$** Cell pellets were thawed on ice, weighed and resuspended in solubilisation buffer (1% n-Dodecyl β-D-maltoside (DDM) (w/v), 20 mM HEPES, 10% (v/v) glycerol, 150 mM NaCl, pH 7.5) at a ratio of 1:10 (w/v) of cell pellet to solubilisation buffer. Pellets were solubilised for 1h on a DigiRoll 6 roller (SLS, UK) at 80RPM and 4°C. Samples were clarified by centrifugation at 4122 x *g* for 20 min at 4°C. Purification of TS-SNAP-$A_{2A}$ was achieved by the use of MagStrep "type3" XT magnetic beads (IBA, Göttingen, Germany). Beads were prepared by removal of supernatant using a magnetic separator (IBA, Göttingen, Germany) and then they were washed twice in solubilisation buffer before being added to samples. Samples were incubated with beads overnight on a DigiRoll 6 roller set to 80 RPM at 4°C. The following morning supernatant was removed from beads using the magnetic separator and beads were washed twice with wash buffer (0.1% DDM (w/v), 10% glycerol (v/v), 150mM NaCl, pH 7.5), before resuspension in 30μL elution buffer (1:9 solution of 10X buffer BXT (IBA) and wash buffer). Elution took place for 5 hours on a DigiRoll 6 roller set to 80 RPM at 4°C. Samples were then separated from beads using magnetic separator and then immediately processed for electrophoresis.

**Gel electrophoresis and in gel fluorescence** 15μL of samples containing purified TS-SNAP-$A_{2A}$ were mixed with 5μL NuPAGE™ LDS sample buffer and resolved on a NuPage™ 4-12% Bis-Tris 15 x 1.0mm well gel using NuPage™ MOPs SDS running buffer. Gels were run for 50 min at 200V. Samples were not boiled prior to gel electrophoresis. 5μL PageRuler™ Prestained Protein Ladder was used as the ladder. Gels were scanned on an Amersham Typhoon imaging system (GE Healthcare Life Sciences, Pittsburgh, PA) using Fluorstage and Cy5 670BP30 filter sets with PMT set to auto and pixel size to 200 μm.

**CRE-SPAP gene transcription.** $A_{2A}$R CRE-SPAP cells were grown to confluence in clear 96-well plates and on the day prior to analysis normal growth medium was removed and replaced with serum-free medium (SFM; DMEM/F12 supplemented with 2 mM L-glutamine) with or without 1 μM **1**. On the day of the experiment, all media was removed from wells, fresh SFM media added and for wells requiring the continued presence of **1**, this media contained fresh 1 μM **1**. Cells were incubated for 30 min at 37°C, and then for the **1** overnight treated wells, SFM was removed and replaced with fresh SFM. Then increasing concentrations of CGS21680 was added to the required well and plates incubated for 5 h at 37°C, 5% $CO_2$. After the 5 h incubation, all medium was removed from the cells and replaced with 40 μl of SFM and incubated for a further 1 h at 37°C. The plates were then incubated at 65°C for 30 min to destroy the endgoneous alkaline phosphatases. After cooling the plates to room temperature, 5 mM 4-nitrophenyl phosphate in a diethanolamine-containing buffer (10% (v/v) diethanolamine, 280 mM NaCl, 500 μM $MgCl_2$, pH 9.85) was added to each well. Plates were incubated for 15 min at 37°C and then the absorbance at 405 nm was measured using a Dynex MRX plate reader (Chelmsford, MA, USA). After fluorescence was measured, gels were stained with InstantBlue® protein stain (Expedeon, Cambridge, UK) overnight. Gels were washed twice with $dH_2O$ for 5 min before visualising using a standard smart phone camera.

**Confocal Microscopy** All confocal microscopy was performed on cells grown on 8-well Labtek chambered coverglasses (Nunc Nalgene). Where appropriate, cells were labelled with 100 nM SNAP-surface Alexa Fluor 488 (SNAP-AF488; New England Biolabs) for 30 min at 37°C/5% $CO_2$ in normal growth media. Cells were washed twice in media prior to the addition of the required compounds (250 nM **1** or 250nM **1**+ 10 μM ZM241385) in normal growth media and incubated for 2h at 37°C/5%$CO_2$ and then imaged. Where required 10 μM ZM241385 was added after 2h and cells incubated for a further 1 h prior to imaging. For wash experiments, cells were imaged after 2h and then washed every 15 min for 1h in normal growth media prior to imaging again. SK-BR3 cells were incubated with or without 10 μM ZM241385 in normal growth media for 30 min prior to the addition of 250 nM **1** and incubated for 2h at 37°C/5% prior to imaging. All imaging was performed using a Zeiss LSM880 confocal microscope (Carl Zeiss GmbH, Jena,

Germany) using a 63x plan-Apochromat NA 1.4 Ph3 oil-immersion objective and a 488/561/633 main dichoroic. For SNAP-AF488 a 488 nm argon laser was used to excite the fluorophore and the emission was detected between 493 and 628 nm. Cy5 was excited using a 633 nm HeNe laser and emission collected between 638 and 747 nm for HEKG cells and between 633 and 695nm for SK-BR3 cells. Due to the lower level of expression in the SK-BR3 cells, a pixel dwell time of 4.12 µs was used, whereas a pixel dwell time of 2.06 µs was used with HEKG cells. Within each experiment, a pinhole diameter of 1 Airy unit was used and the laser power, gain and offset kept constant for each experiment set. To obtain membrane intensity values, regions of interest were drawn by hand around the cell membranes in Zen Black image analysis software.

**Data analysis**

We simultaneously fit the total and nonspecific saturation binding curves using the following equation:

$$TRFRET\ ratio = \frac{B_{max} \times [B]}{[B] + K_D} + \left((M \times [B]) + C\right)$$

where $B_{max}$ is the maximal signal, [B] is the concentration of fluorescent ligand in nM, $K_D$ is the equilibrium dissociation constant in nM, M.is the slope of the non-specific binding component and *C* is the intercept with the y-axis.

The CRE-SPAP data were fit to the following equation:

$$Response = \frac{E_{max}.[A]}{[A] + EC_{50}}$$

where $E_{max}$ is the maximal response, [A] is the concentration of agonist and the $EC_{50}$ is the molar concentration of agonist required to generate 50% of the $E_{max}$

We carried out statistical analysis using two-tailed paired or unpaired t test as required.

### Supplementary Figure Legends

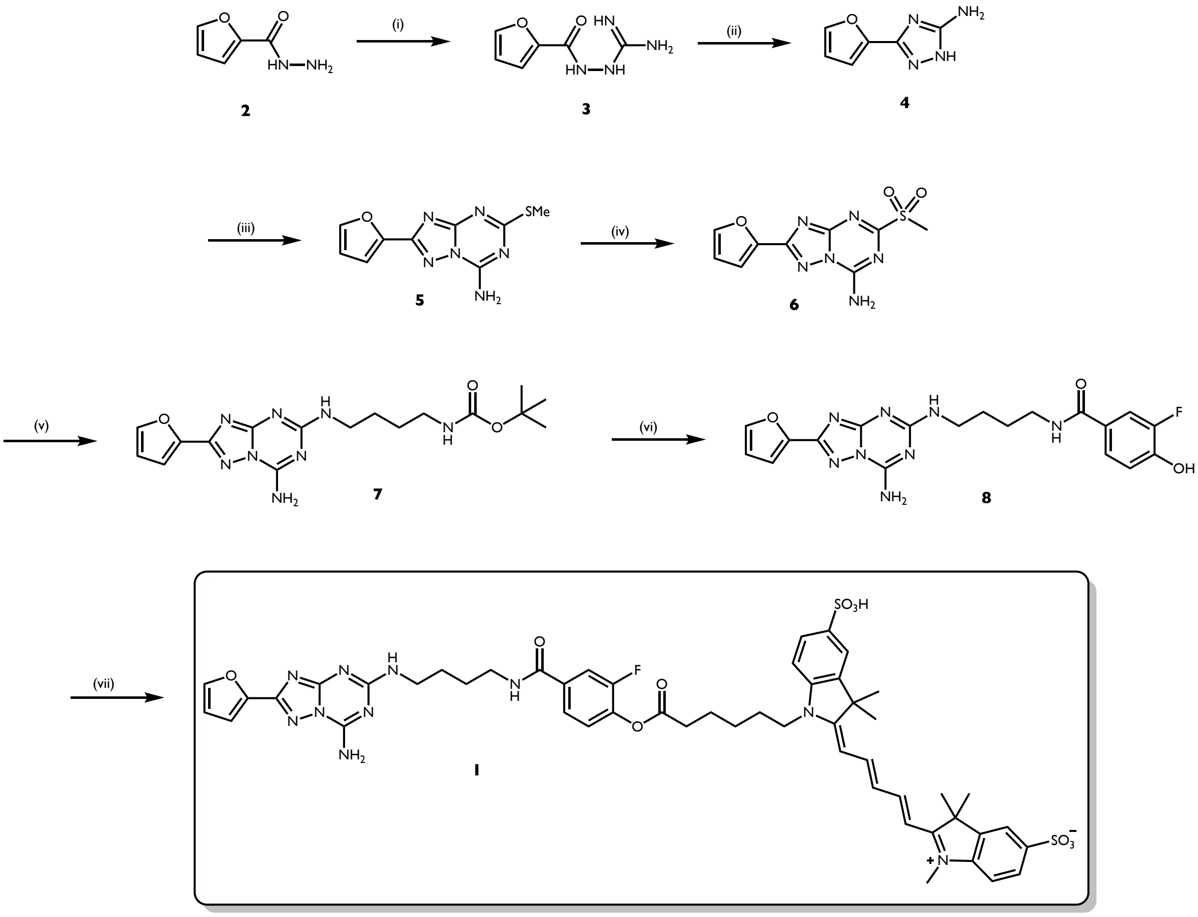


**Supplementary Figure 1. Synthesis Scheme for Fluorescent Probe 1**

*Reagents and Conditions:* (i) 2-methyl-2-thiopseudourea hemisulfate salt, $NaOH_{(aq)}$, 24 h; (ii) $H_2O$, **Δ**, 24 h; (iii) Dimethyl *N*-cyanodithioiminocarbonate, 170 ℃, 1 h; (iv) 3-chloroperbenzoic acid, dichloromethane, 16 h; (v) *tert*-Butyl *N*-(3-aminobutyl)-carbamate, acetonitrile, 2 h; (vi) (a) Trifluoracetic acid, dichloromethane, 0.5 h; (b) 3-fluoro-4-hydroxybenzoic acid, 1-[Bis(dimethylamino)methylene]-1*H*-1,2,3-triazolo[4,5-*b*]pyridinium 3-oxide hexafluorophosphate, diisopropylethylamine, *N,N*-dimethylformamide, 90 ℃, 2 h; (vii) sulfo-cyanine-C5 carboxylic acid, 2-Bromo-1-ethyl-pyridinium tetrafluoroborate, diisopropylethylamine, *N,N*-dimethylformamide, 16 h in the dark.

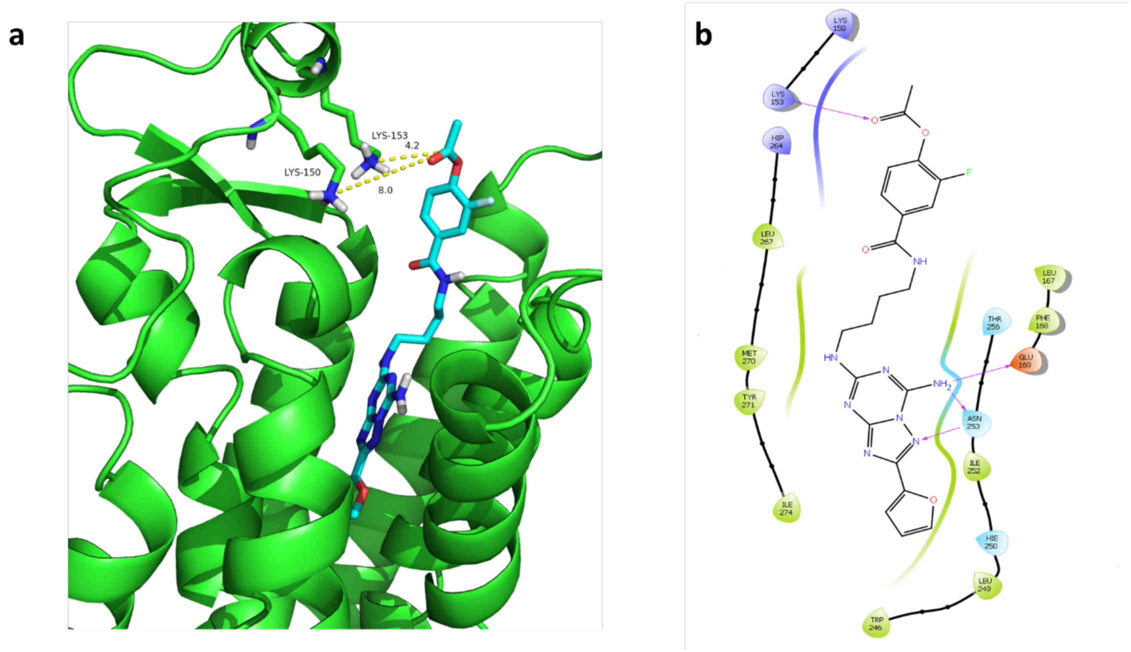


**Supplementary Figure 2. Representation of the binding mode of truncated (1) in the human $A_{2A}$ Receptor ($hA_{2a}R$).** a) Molecular docking pose of truncated (**1**) (Cyan) in the $hA_{2A}R$ (green ribbon, PDB: 4EIY); b) 2D ligand-protein interaction diagram for truncated (**1**) with key $hA_{2A}R$ binding site residues, purple arrows indicate hydrogen bonds interactions.

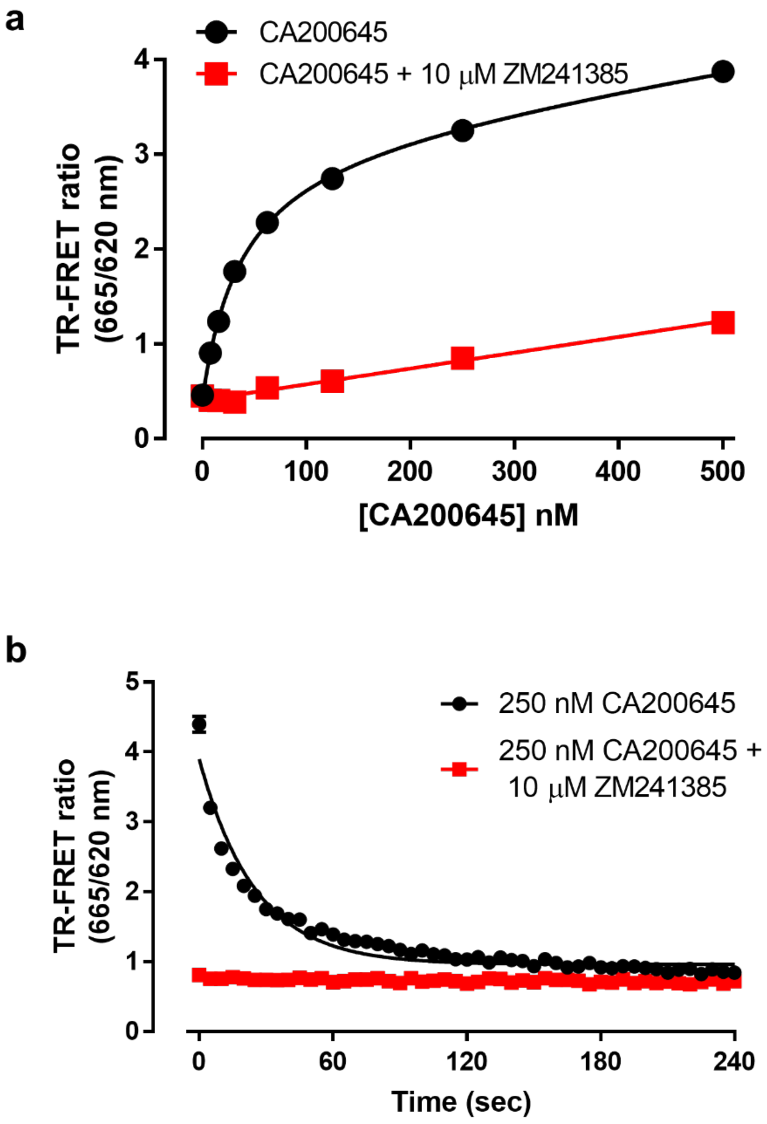


**Supplementary Figure 3. Pharmacological characterisation of CA200645 at SNAP-$A_{2A}$** (**a**) TR-FRET saturation binding curves obtained by treating membranes containing Lumi4-Tb labelled SNAP-$A_{2A}$R with increasing concentrations of CA200645 in the absence (black circles) or presence (red squares) of 10 μM ZM241385 for 1h at 37$^{o}$C prior to determination of TR-FRET ratio. Data shown is representative of five experiments and each data point represents mean ± s.e.m of triplicate determinations. Calculated $K_d$ for CA200645 at SNAP-$A_{2A}$ was 36.8 ± 5.7 nM.(**b**) Membranes containing Lumi4-Tb labelled SNAP-$A_{2A}$R were pre-treated with 250 nM CA200645 in the presence (red squares) or absence (black circles) of 10 μM ZM241385 for 2h green squares at 37$^{o}$C prior to measurement of TR-FRET ratio. After basal reads, 10 μM ZM241385 was added to all wells and measurements taken every 5 sec for 5 min.  Data shown is representative of four experiments and each data point represents mean ± s.e.m of triplicate measurements.

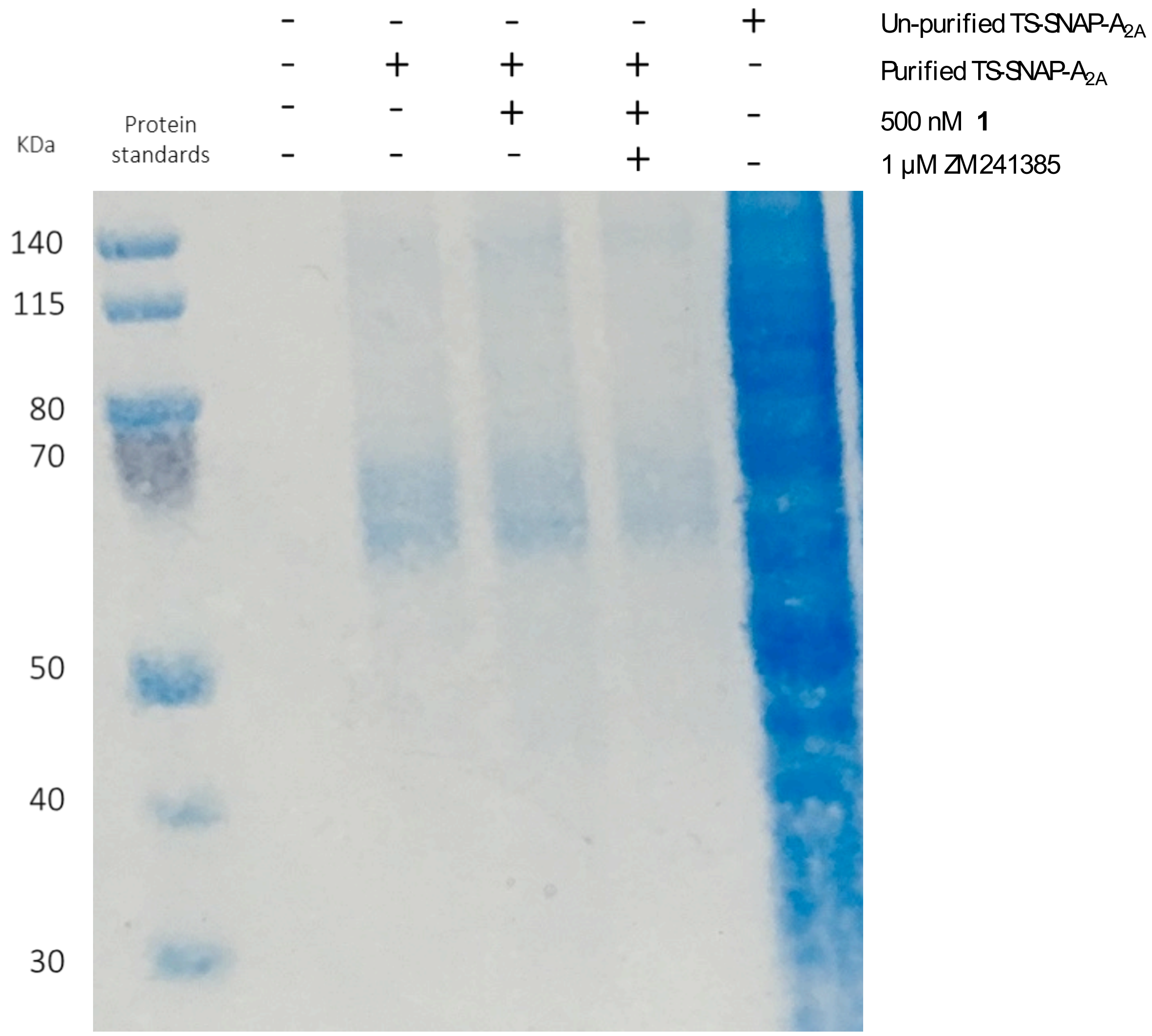


**Supporting Figure 4. Protein stain of 1 labelled samples**

T-Rex™-293 cells induced to express TS-SNAP-$A_{2A}$R were treated with 500nM **1** in the presence or absence of 1μM ZM241385. Crude cell pellet extract from untreated cells before and after purification were used as controls. Where indicated TS-SNAP-$A_{2A}$R was purified and all samples were analysed on an SDS-PAGE gel. Direct Cy5 fluorescence was visualised using in-gel fluorescence and shown in Figure 1e. After in-gel fluorescence was visualised, gel was stained for using InstantBlue® protein stain to demonstrate equal loading and purification and image obtained on a standard smartphone camera. Gel shown is representative of three independent experiments.

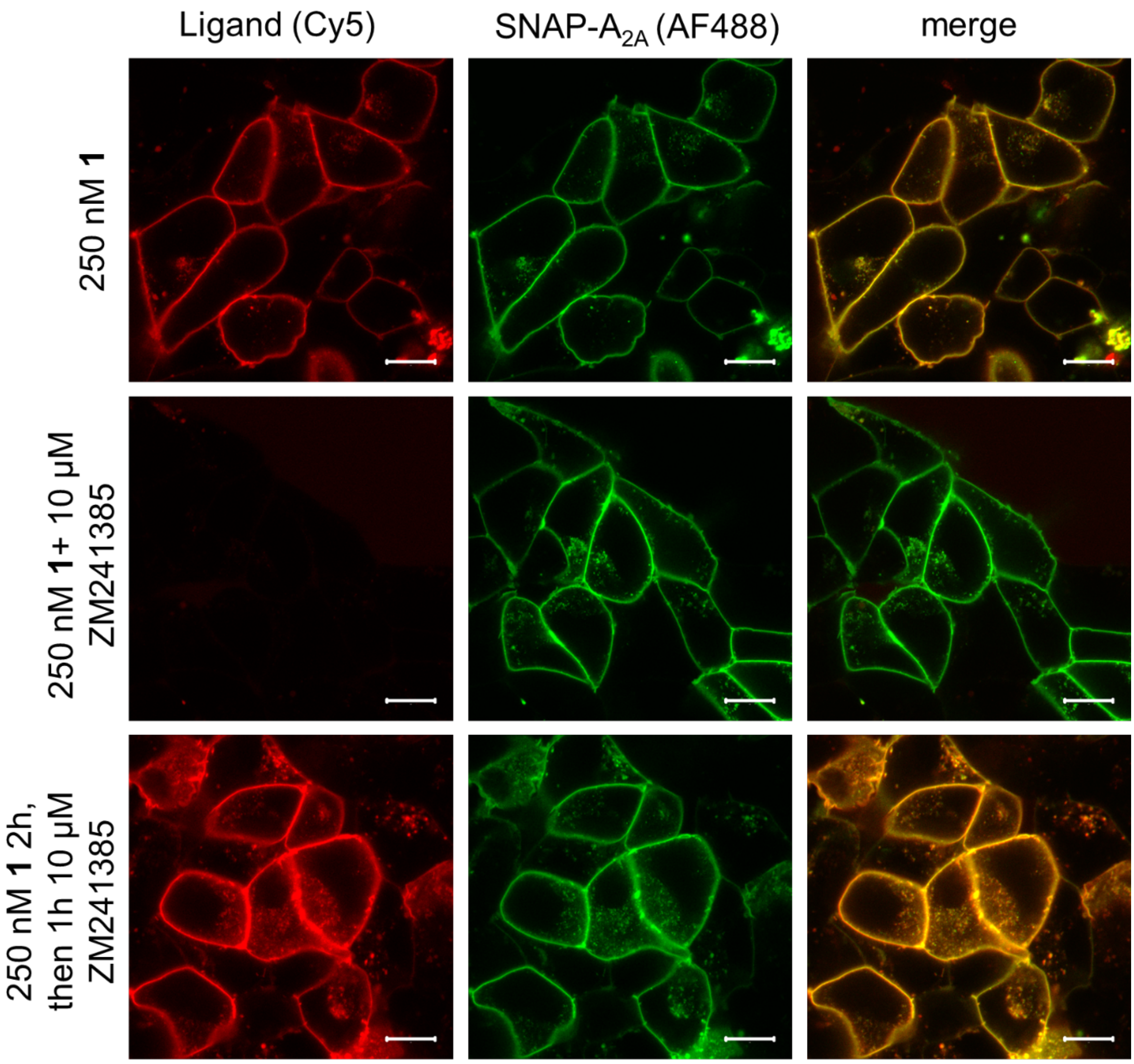


**Supplementary Figure 5. Visualisation of Cy5 labelling with 1 after treatment with ZM241385** Live HEK293 cells expressing SNAP-$A_{2A}$R were labelled with SNAP-surface-AF488 and then treated with 250 nM **1** in the absence (top row) or presence (middle row) of ZM241385 for 2h prior to the capture of single equatorial confocal images. To the cells treated with 250 nM **1**, 10 µM ZM241385 was added and incubated for a further 1h then imaged (bottom row). Left hand column represents Cy5 fluorescence, middle column AF488 fluorescence and right hand column the merged image. Images shown are representative of images taken in four independent experiments, with all image taken using identical settings for laser power, gain, and offset in both channels. Scale bar shown equals 10 µm.

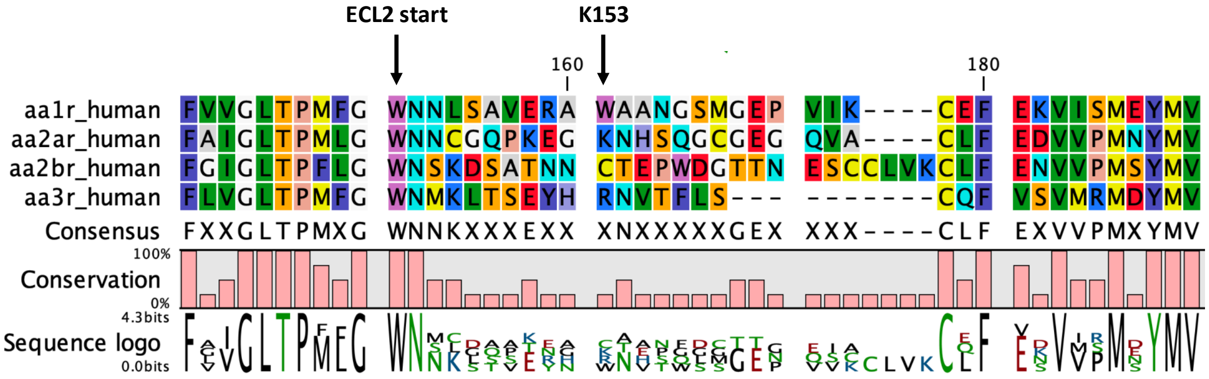


**Supplementary Figure 6 Sequence alignment of extra cellular loop 2 of the four adenosine receptor subtypes**

Alignment of the top of transmembrane region 5, extra-cellular loop 2 (ECL2), and top of transmembrane region 6 of the four adenosine receptor subtypes. The potential site of Cy5 attachment to K153, by **1** in the $A_{2A}R$ is indicated by an arrow which is not conserved in the other three adenosine receptor subtypes.

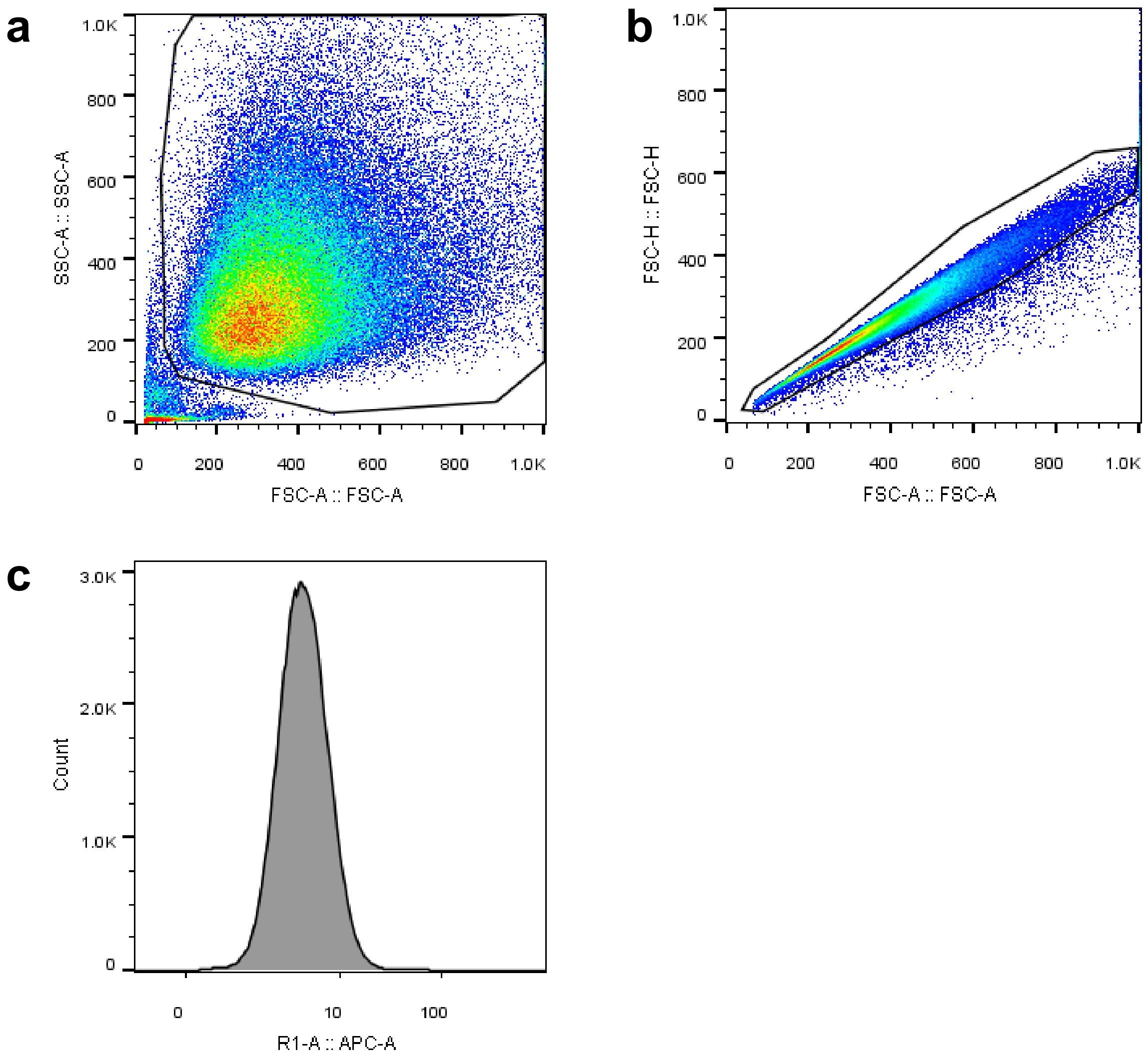


**Supplementary figure 7 FACS gating strategy for macrophages**

Macrophages were harvested on ice after incubating with **1** in the presence or absence of 10 µM ZM241385. AFSC/SSC plot was used to gate out debris (a), followed by Doublet exclusion (b) and singlet macrophages plotted as a histogram in R1 channel ('APC') for Cy5 fluorescence from NDK174 (c). Example shown is a representative sample (**1** + 10 µM ZM241385) from a representative donor (one donor of four).
